# Pancreatic tumor organoids for modeling in vivo drug response and discovering clinically-actionable biomarkers

**DOI:** 10.1101/513267

**Authors:** Ling Huang, Bruno Bockorny, Indranil Paul, Dipikaa Akshinthala, Omar Gandarilla, Arindam Bose, Veronica Sanchez-Gonzalez, Emily E Rouse, Sylvain D. Lehoux, Nicole Pandell, John G. Clohessy, Joseph Grossman, Raul Gonzalez, Sofia Perea Del Pino, George Daaboul, Mandeep S. Sawhney, Steven D. Freedman, Richard D Cummings, Andrew Emili, Lakshmi B Muthuswamy, Manuel Hidalgo, and Senthil K Muthuswamy

## Abstract

Patient-derived models are transforming translational cancer research. It is not clear if the emergence of patient-derived organoid (PDO) models can extend the utility of the widely used patient-derived xenograft (PDX). In addition, the utility of PDO models for serum biomarker discovery is not known. Here, we demonstrate that PDO models recapitulate the genomics, cell biology, glycomics and drug responses observed in PDX models. Furthermore, we demonstrate the applicability of PDO models for identification of N-glycans that are enriched in the glycome of pancreatic ductal adenocarcinoma (PDAC). Surprisingly, among all the glycans observed in PDX and PDOs, a core set of 57 N-glycans represent 50-94% of the relative abundance of all N-glycans detected, suggesting that only a subset of glycans dominate the cell surface landscape in PDAC. In addition, we outline a tumor organoid-based pipeline to identify surface proteins in extracellular vesicles (EV) from media supernatant of PDO cultures. When combined with the affinity-based validation platform, the EV surface proteins discovered in PDOs are effective in differentiating patients with PDAC from those with benign pancreatitis in the clinic, identifying PDO as powerful discovery platform for serum biomarkers. Thus, PDOs extend the utility of the archival collections of PDX models for translational research and function as a powerful platform for identification of clinically-actionable biomarkers in patients blood.

**Significance statement:** Tumor organoids extend the utility of PDX models as platforms for investigating drug response, glycosylation changes and function as new platforms for discovering blood-based biomarkers

---

Translational cancer research has benefited significantly from the use of patient materials either for genomics analysis or for generation of patient tumor derived models such as xenograft (PDX) or organoid cultures. These patient derived models frequently retain inter- and intra-patient variations of the disease, which is a significant advantage over the traditionally used immortalized cancer cell lines that suffer from genetic drift and variance from original patient tumors due to long-term maintenance in culture. Over the years, a great deal of efforts has been placed on generation of PDX models representing most cancer types including pancreatic ductal adenocarcinoma (PDAC). PDX models are known to not only maintain multiple aspects of cancer traits, but also model therapeutic drug response in patients with more than 85% accuracy (1). Despite their high fidelity in modeling human cancer, routine use of PDX models for all aspects of translational research is impeded by their high cost and the extended time needed to conduct experiments. Furthermore, PDX models are not an ideal platform for large scale studies aimed at screening multiple therapeutic drugs and their combinations due to high cost and time. Patient tumor derived organoid (PDO) models are fast evolving as a method to model a patient’s disease *ex vivo*, such as studies in pancreatic cancer (2, 3), however, it is not clear if PDO models phenocopy PDX models in translational cancer research. In particular, it remains to be determined whether PDOs can serve as a discovery platform that can be used in conjunction with PDX models to discover clinically relevant findings.

Apart from therapies, identifying new clinically relevant blood-based biomarkers is a pressing need for diagnosing cancer and for monitoring disease (4–6). In particular, identifying blood-based biomarkers, such as extracellular vesicle associated proteins, that can distinguish cancer from benign diseases is an extremely challenging task. Using PDX to discover new secreted vesicle-based biomarkers is a non-trivial task in part due to presence of contaminating host factors. Whether PDO cultures, that are made-up of cancer cells, can be used to identify new biomarkers has not been explored.

Beyond drug response and genomics, glycosylation have important implications for therapeutics and for biomarker identification. Both N-glycosylation and O-glycosylation are significantly altered in malignant tissues and such changes can profoundly impact protein function in multiple ways, including protein maturation, localization, folding, cell adhesion and trafficking, cell signaling, and immune response (7). However, glycosylation changes are rarely studied in patient derived models, likely due to resource limitations associated with using PDX model. Here, we investigated if PDO cultures will serve as a platform to identify changes in glycosylation that are relevant *in vivo*.

To demonstrate the power of using PDO models for translational research as a whole, including identification of clinically relevant biomarkers, we conducted prospective studies to demonstrate that PDOs are effective in predicting drug responses observed *in vivo* PDX models; effective in identifying common glycosylation features enriched in PDAC tumors and are effective models for identifying secreted extracellular vesicles (EV) protein markers that differentiate patients with PDAC from those who have chronic pancreatitis.

## Results

### Tumor organoids recapitulate genomic profiles of matched PDX tumors

To build a set of matched PDX and tumor organoid models, we identified eight PDX models that have heterogeneous combination of alterations in the common pancreatic cancer oncogenes such as KRAS, TP53, CDKN2A, SMAD4, cMyc, GATA6, ERBB2, as well as genetic alterations in SWI/SNF, DNA damage repair and axon guidance pathways (8–10) (**Fig.1A**) as determined by exome sequence analysis of the PDX tumors. The models include one KRAS^wt^, one KRAS^G12V^, one KRAS^Q61H^, and five KRAS^G12D^ (**Fig.1A**). All models are PDAC except Panc014, which is cholangiocarcinoma (see **Fig.S1A** for patient information). We generated organoids from those models using a method described previously (11). Comparison of exome sequences of six organoids and their matched PDX models revealed overall conservation of genomic alterations, with the exception of one allele Notch2 (**Fig.1B**). Twenty-four of the twenty-five allelic alterations of frequently mutated genes in PDAC matched between tumor organoids and PDX tumors, demonstrating that organoid culture maintained critical genomic features of the matched PDX tumors.

To determine if tumor organoids recapitulate biology and architecture of the PDX models, we analyzed organoid sections by hematoxylin & eosin stain and using a molecular marker (cytokeratin 19) that is expressed in ductal epithelia of the pancreas. Panc014 PDX tumors had glands predominantly cribriform in architecture, with focal poorly differentiated areas; Panc030 PDX tumors had glands that were confluent, with very focal single cells; Panc281 PDX tumors had glands somewhat cribriform/confluent but still easily identifiable as glands, with focal poorly differentiated areas (**Fig.1C).** Tumor glands were consistently observed in all organoid cultures. Overall, the correlation of histopathological features between PDX and organoids was very apparent in Panc014 and Panc281, while in Panc030 it was less clear, demonstrating that organoid cultures mostly retain inter-model variations in histopathology. Tumor cells in organoids and PDX tumors expressed cytokeratin 19 (*panel (II)* in **Fig. 1C**) demonstrating that cells in culture and PDX models retained pancreatic epithelial differentiation.

**Figure 1.**
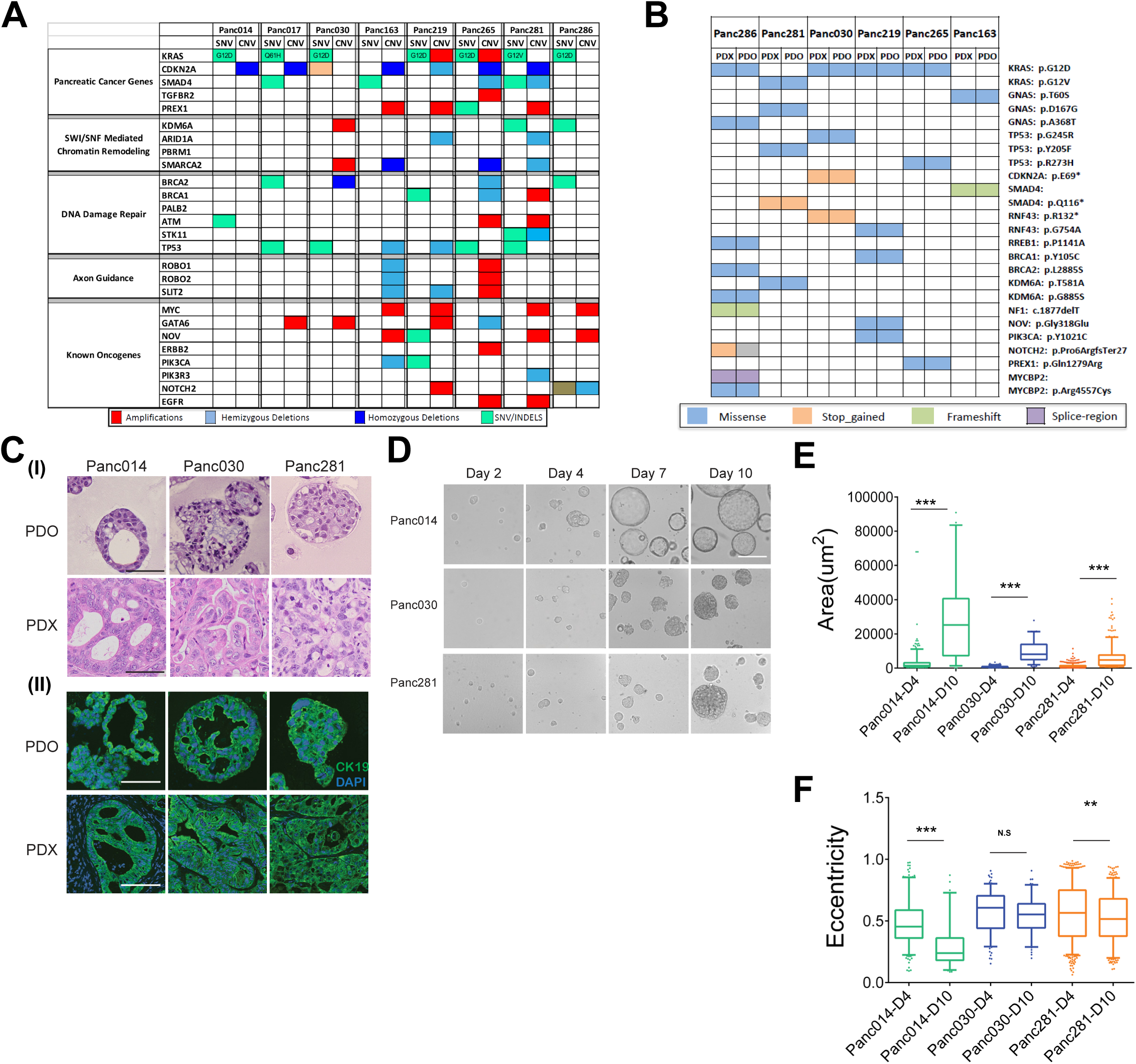
Genomic and glycomic profiles of tumor organoids and matched PDX tumors **(A)** Genomic pathways associated with PDAC observed in PDX models. **(B)** Concordance of mutant alleles between tumor organoids and matched PDX tumors. **(C)** H&E and immunofluorescence images of tumor organoids and matched PDX tumors. Panel (I), H&E images; panel (II), immunofluorescence images, DAPI, blue, cytokeratin 19, green. Scale bars, 50 µm. **(D)** Phase contrast images of organoids at different stages of cultures. Scale bars, 50 µm. **(E)** Area distribution of organoids derived from Panc014, Panc030 and Panc281 at day 4 and day 10. **(F)** Eccentricity distribution of organoids derived from Panc014, Panc030 and Panc281 at day 4 and day 10.

Previous studies have demonstrated a relationship between phase contrast morphology of cells in 3D culture and their gene expression profiles (12), suggesting that differences in organoid morphology can provide important insights into tumor biology. Organoids from three tumor models with different levels of genomic alterations, Panc014, Panc030 and Panc281, (**Fig.1D**) demonstrated high diversity in organoid morphology as observed by wide range in area within and between different organoids lines suggesting the maintenance of inter- and intra-tumor heterogeneity in organoid cultures. At day 10, organoids from Panc014 were cystic and had smooth edges while Panc030 and Panc281 were solid coinciding with lower level of genomic alterations observed in Panc014 **(Fig.S1A)**. To better understand differences in other morphometric parameters, we used OrganoSeg, a segmentation and analysis platform that can extract multiple parameters from spheroid and organoid cultures (13). As shown in **Fig.S1B**, individual organoids were identified and highlighted based on their phase contrast intensity profiles to create a mask and measure the area (as index for organoid sizes) and eccentricity (as index for organoid circularity). An organoid with lower eccentricity is closer to a perfect circle (eccentricity for a perfect circle is 0). In all three lines of organoids, areas increased significantly between day 4 and day 10 in culture (**Fig.1E**). We also confirmed this by measuring cell proliferation using CellTiter-Glo 3D. Interestingly, Panc014 organoids displayed a linear increase in cell number compared to Panc030 and Panc281 despite having low mutation burden (**Fig.S1A,C**). This was also consistent with their difference in growth potential *in vivo* (**Fig.S1D**). Organoids in Panc014 and Panc281 also became more circular from day 4 to day 10 during growth in 3D while changes in Panc030 (**Fig.1F**) were not significant. These observations highlight the potential for carefully monitoring and analyzing differences in phase contrast morphology of organoid cultures.

### Tumor organoids predict response to drugs *in vivo*

Response to therapeutic drugs in PDX models is known to correlate well with patient response to the same therapies (1). To determine whether organoids can serve as a complement to PDX models, we investigated if organoids can phenocopy drug responses in PDX models. We reasoned that a successful outcome will help demonstrate the utility of the organoid models as a cost effective and scalable platform for preclinical and translational research efforts aimed at finding new drugs and drug combinations using patient-derived models.

We tested four drug combinations, two representing standard of care for PDAC: gemcitabine/paclitaxel (to model gemcitabine/nab-paclitaxel) and 5-fluorouracil (5FU)/oxaliplatin; and two combination therapies studied in clinical trials: gemcitabine/olaparib and paclitaxel/palbociclib. To best model tumor’s exposure to drugs *in vivo*, we chose physiologically permissive drug concentration ranges using reported peak plasma concentrations (C_max_) as a reference point for each drug (14–17). The only exception was 5FU, which is known to have a half-life of 8 minutes *in vivo*, therefore, we set the peak concentration for 5FU to be 10 µM which was approximately 10 fold over the steady state concentration (18). In addition, the drug combinations were chosen in reference to the clinically meaningful ratio of the two drugs. For gemcitabine/paclitaxel, 5FU/oxaliplatin, and gemcitabine/olaparib, we started with C_max_. For paclitaxel/palbociclib, our tested concentrations started from 3-fold of C_max_ since this is an investigational combination for PDAC (**Fig. 2A**). As shown in **Fig.2B**, all four combinations induced dose dependent growth inhibitions in organoids as determined by cell viability measurements, though the degree of effect varied. To compare effects among treatments within a sample, we calculated the area under the curve (AUC) of each treatment then normalized to AUC of the least effective treatment. For example, for Panc163 organoids, 5FU/oxaliplatin (red bar) was least effective while gemcitabine/paclitaxel (blue bar) had highest inhibitory effects. To test whether differences in organoid drug responses related to differences in drug sensitivities *in vivo*, we tested gemcitabine/paclitaxel and 5FU/oxaliplatin treatments in Panc163 PDX models (for dose and schedule, see **Fig.S3**). As shown in bottom panel of **Fig.2B**, gemcitabine/paclitaxel (blue line) was more effective than 5FU/oxaliplatin (red line) in suppressing tumor growth *in vivo*. In addition to Panc163, differential sensitivity to drugs in Panc014, Panc030, Panc281 organoids correlated to their responses *in vivo* (**Fig.2C**). In Panc014, gemcitabine/paclitaxel and gemcitabine/olaparib treatments induced similar tumor suppression both in PDX and PDO models. Interestingly, in all organoids tested, paclitaxel/palbociclib treatments were inferior to gemcitabine-based regimens suggesting gemcitabine needed to be added to the combination to achieve better effects.

**Figure 2.**
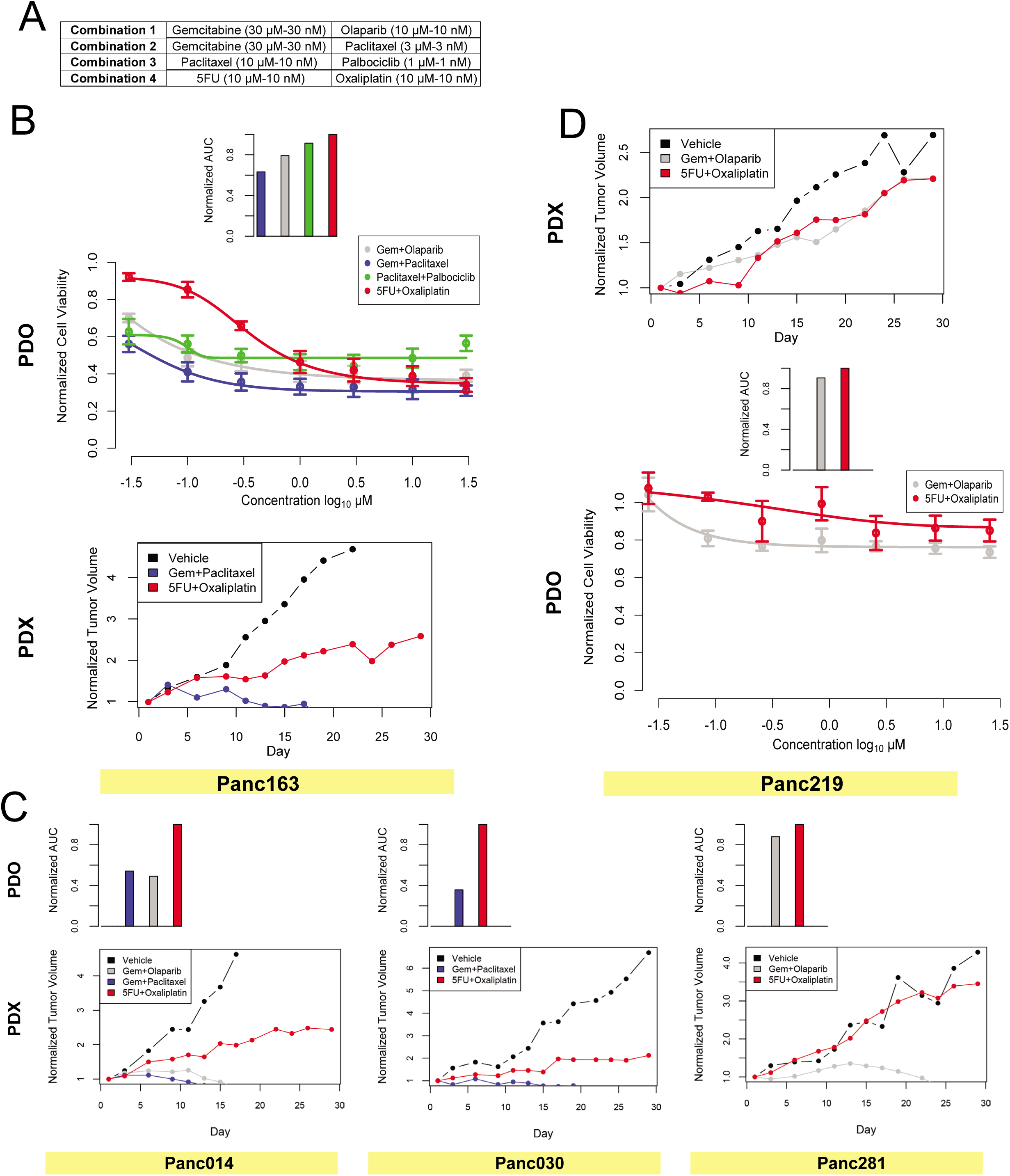
Comparison of drug responses between tumor organoids and matched PDX models **(A)** Dose ranges of drug combinations tested in tumor organoids. **(B)** Bar chart represents normalized area under the curve (AUC) values calculated by measuring changes in cell viability of Panc163 organoids treated with seven increasing doses of respective drug combinations. Middle graph represents changes in cell viability in response to drug treatments of Panc163, which is used to prepare the AUC charts. Bottom panel represents changes in tumor volume of Panc163 PDX tumors treated with indicated drug combinations. **(C)** Top panel represent normalized AUC values indicating changes in cell viability of organoids treated with seven increasing doses of the indicated drugs. Bottom panel represents changes in tumor volume of PDX tumors treated with indicated drug combinations. **(D)** Top panel represents changes in tumor volume of Panc219 PDX tumors treated with indicated drug combinations. Bottom panel represents AUC and changes in cell viability of organoids treated with increasing concentrations of the indicated drugs.

**Figure 3.**
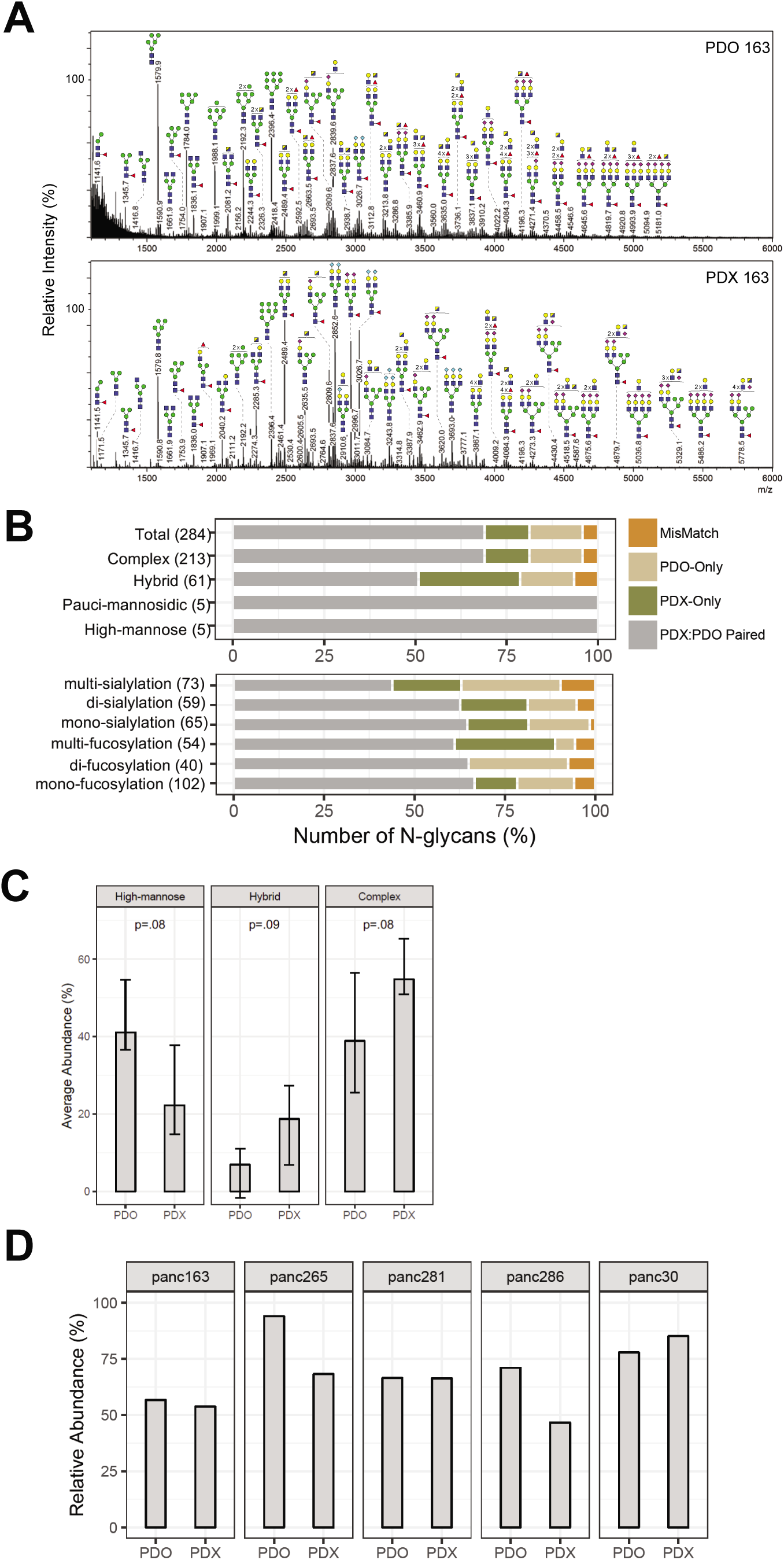
Glycomic analysis on organoids and matched PDX models **(A)** Representative mass spectrometric N-glycans profile in PDO and matched PDX from Panc163. Cartoons depict glycan composition for respective m/z peaks. **(B)** N-glycan classes and their occurrence in PDO and matched PDX models. Numbers in Y axis represent total N-glycan in each class and X-axis shows percentage of occurrence in PDX and PDO models as indicated by the color charts **(C)** Average relative abundance of three major N-glycan classes in PDX and PDO models. **(D)** Total relative abundance of 57 common N-glycans for each model.

In our studies with PDX models, we identified Panc219 as a model that was resistant to both gemcitabine/olaparib and 5FU/oxaliplatin *in vivo* (top panel, **Fig.2D**). To investigate if resistance to treatment *in vivo* is also observed in organoid cultures, we generated organoid models for Panc219 and investigated the drug combinations. Interestingly, organoids were resistant to both regiments with less than 20% decrease in cell viability even at the highest drug concentrations tested (bottom panel, **Fig.2D**) demonstrating the ability of organoid platform to model *in vivo* drug responses. These findings strongly suggest that organoid platform could complement patient derived xenograft models for investigating drug sensitivities and serve as a scalable, discovery platform in translational research efforts using existing and new PDX models.

### Tumor organoids identify a conserved set of glycans in PDAC

Like the genome of cancer cells, glycosylation of cellular proteins is known to undergo significant and varying alteration in cancer cells (19). Changes in glycosylation will have a dramatic impact on protein function including its expression levels, stability, and localization (7). N-glycan and O-glycan profiles are known to vary among pancreatic cancer cells (20). Since glycosylated proteins are widely used as biomarkers and as therapeutic targets (21), developing a better understanding of glycosylation changes using patient derived models will be of significant advantage to the field.

To determine if organoids can serve as models to understand glycosylation in patient derived tumor models, we compared the N-glycan profiles of matched PDX and PDO models. Five pairs of PDX and PDOs were lyophilized, digested with trypsin then with PNGaseF to release N-glycans and the permethylated glycans were analyzed by MALDI-TOF. Representative N-glycan profiles for a PDX-PDO pair is shown in **Fig.3A and Fig.S3A** with a few glycan masses annotated to demonstrate the overall similarity in spectrum between PDX and PDO samples. By analyzing 10 samples (5 PDX and 5 PDO), we identified a total of 284 N-glycan masses and predicted structures. Interestingly 66% (188) of them were shared by both PDX and PDO (**Fig.3B**).

To determine if there are differences between PDX and PDO models in the subtypes of glycan modification, we first classified N-glycans into four major types: a) high-mannose – five to nine mannose moieties are attached to the core, b) pauci-mannose – less than three or less mannose are attached to the core, c) complex: where ‘antennae’ are initiated by N-acetylglucosamine, and d) hybrid: where mannose residues and one or two antennae are attached to the core (7). Amongst the total number of N-glycans (284) identified in all models, complex N-glycans represented 87-90% of these, followed by hybrid and high-mannose types. In terms of concordance between N-glycans observed in PDX and PDO models, high-mannose showed 100% and complex type of glycan showed 74%-90% concordance in terms of occurrence, demonstrating that the subtypes do not differ between PDX and PDO models. Occurrence of complex N-glycans with and without fucosylation and/or sialylation also showed high overlap between PDX and PDO further demonstrating that the PDOs retain the glycan diversity observed in *in vivo* contexts (**Fig.3B**).

We next compared the relative abundance of three major N-glycan types to rule out the possibility that they differ between in PDX and PDO samples (**Fig.3C**). On an average, the complex type was most abundant in both PDX and PDO models, followed by high-mannose and hybrid subtypes. Thus, in addition to similarities in occurrence of N-glycan types, quantitative comparison of relative abundance of different N-glycans showed high concordance between PDX and PDO models.

Our study also provided an opportunity to investigate the diversity and commonality of glycan among all the samples analyzed. When glycan profiles were analyzed in samples individually and collectively, we observed that PDX models on average have 143+/-21 glycans and PDO models on an average have 138+/-8.9 glycans demonstrating that not all 284 glycans are observed in every sample analyzed. Fifty-seven N-glycans were present in all 10 samples. Interestingly, these 57 N-glycans (5-high mannose, 3-pauci-mannose, 7-hybrid and 42-complex) together represent 53% to 94% of total N-glycans observed in these samples **(Fig.3D**), demonstrating that these 57 N-glycan masses dominate the N-glycans landscape in PDAC samples. This unexpected observation raises the possibility that PDAC samples may have shared glycosylation signature and a shared underlying mechanism that drives these glycosylation patterns.

### Organoids as discovery platform for blood-based biomarkers in PDAC patients

There is a significant need for diagnostic biomarkers in PDAC. This is a highly active field with several lines of investigation in progress. One such promising approach involves identification of secreted extracellular vesicles (EV) in the blood of patient to differentiate patients with PDAC from both patients who are disease-free or those with chronic pancreatitis (benign conditions). Several studies have attempted to identify EV associated protein using conventional cell lines in culture with further validation using blood samples from PDAC patients and healthy controls (4, 22–24). It is not clear whether organoid cultures can be used to discover new, clinically significant, secreted biomarkers. Typically, EV identification effort using cells in culture requires large amount of media supernatant (0.1-1.0 liters), which would be a technical challenge when using organoid cultures. To overcome this bottleneck, we optimized a vesicle enrichment method to concentrate vesicles from 4.0 ml of organoid media supernatant and subjected to LC-MS/MS. To differentiate cancer-associated EV from those secreted by normal human pancreatic epithelial, we used media from our human embryonic stem cell derived exocrine pancreas organoids. Among the 1,465 proteins identified, we identified 241 proteins that were at least two-fold higher in tumor organoid EVs compared to exocrine organoids and expressed in at least 4 out of the 6 tumor organoid lines. Principle component analysis showed that the models differ from each other in the EV proteome. Furthermore, we noted that Panc014 was far removed from the other models analyzed **(Fig. 4A).** Interestingly, Panc014 was derived from a cholangiocarcinoma whereas all other models originated from PDAC demonstrating our ability to detect distinct EV proteomic profiles among patient derived models. In addition, it also highlights the fact that the culture/media conditions used to generate and maintain these organoid models do not induce neutralization of phenotypes but are effective in retaining inter-patient heterogeneity in tumor biology. To obtain a broader understanding of the molecules present in EV, we analyzed proteins that were two-fold higher in tumor organoid EVs compared to normal organoids in the majority of PDAC organoid lines. There were 362 proteins identified, that clustered into functional groups including RNA splicing, histone/chromatin, proteasome and translation, in addition to vesicles containing proteins involved in cytoskeleton regulation, cell adhesion and membrane trafficking (. These observations are consistent with previous reports of proteins present in exosomes and other extracellular vehicles, demonstrating the ability of organoid platform to identify patient-specific differences in extracellular vesicles (4, 22–24).

**Figure 4.**
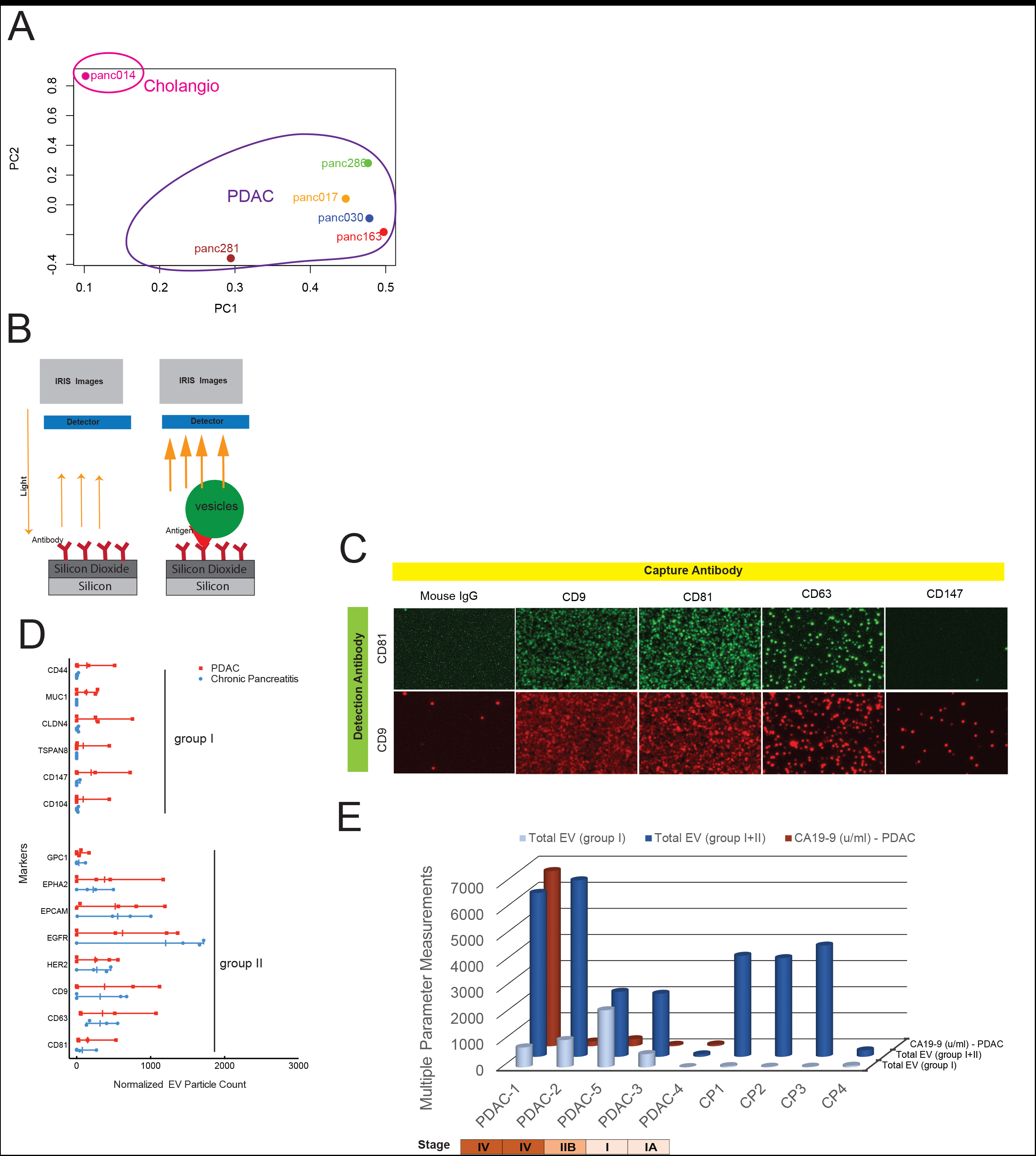
Identification of EV associated proteins in tumor organoid media and validation in patient blood. **(A)** Principle component analysis on proteins abundance in EV present in culture media from tumor organoids. Organoids representing PDAC or cholangiocarcinoma are indicated. **(B)** Functional clustering of EV associated proteins enriched in tumor organoid culture media. **(C)** Scheme of the Exoview detection technology for quantitation of EVs positive for an antigen of interest. **(D)** Immunostaining of EVs captured by the Exoview affinity chip for known vesicle associated markers. **(E)** Quantification of EVs positive for specific markers in blood from patients with chronic pancreatitis or PDAC. **(F)** Measurements of CA19-9 (maroon) in blood from patients with PDAC and quantification of EV positive for any marker (blue) or only group I markers (grey) in blood from patients with chronic pancreatitis or PDAC.

To demonstrate that extracellular vesicles identified in our organoid platform have clinical significance, we investigated if vesicles containing those proteins identified in organoid culture supernatant can be detected in blood samples of pancreatic cancer patients and if they are specific to patients with pancreatic cancer but not in patients with a non-malignant pancreatic disease. We selected a set of surface proteins that included candidates from our study and those reported by others as exosome associated proteins in gastrointestinal cancers; CD44v6, MUC1, TSPAN8, CD147, CD104 GPC1, CLDN4, EPHA2, EPCAM, EGFR, and HER2. In addition, established exosome markers CD63, CD81 and CD9 were included as control. Blood from five PDAC patients representing various disease stages and four chronic pancreatitis patients with no clinical evidence of cancer were obtained from an IRB-approved protocol for this analysis. To enable detection and quantitation of extracellular vesicles in patient blood, we used a novel affinity microarray-based technology coupled with a Reflectance Imaging Sensor detection methodology (Exoview) to detect and count extracellular vesicles with 50 nm resolution (25). Antibodies against the selected proteins were arrayed on the chip and patients’ plasma were analyzed for presence of marker-positive extracellular vesicles (**Fig.4B**). In addition, the vesicles were immunostained for exosome-specific proteins CD81 and CD9 to demonstrate that portion of the circulating vesicles detected in plasma contain exosomal specific proteins. (**Fig.4C**). Interestingly, the EVs containing antigens against CD44, MUC1, CLDN4, TSAPN8, CD147 and CD104 (Group I) were very low or zero in chronic pancreatitis patients whereas up to 100-fold higher in blood of 3 out 4 PDAC patients (**Fig.4D**). In contrast, EV containing antigens against GPC1, EPHA2, EPCAM, EGFR and HER2 (group II) were detectable in all samples and did not show any selectivity for patients with PDAC. Interestingly, five out of six proteins in group I were identified in our study, in contrast, only four (GPC1, EPHA2 CD9 and CD81) out of eight in group II were detected in our EV analysis, demonstrating the ability of our PDO cultures to serve as a discovery platform to identify EV-associated proteins that can serve as blood-based diagnostic biomarkers for patients with PDAC.

Serum concentration of CA19-9 is the clinically approved diagnostic biomarker for monitoring PDAC progression (26). In the patients’ samples analyzed, CA19-9 levels varied dramatically and showed no association with disease stage (**Fig.4E**), as has been reported. Interestingly, the total EV numbers in group I did not show correlation with disease stage despite their high specificity for PDAC patients over pancreatitis patients. Our results demonstrate that detection of specific type of EVs from patients has great potentials as a diagnostic tool to evaluate different clinical parameters and PDO cultures can serve as platforms to optimize the design of EVs based early detection of pancreatic cancer using patient’s blood.

## Discussion

Organoid cultures have garnered significant interests as a research tool in cancer biology. However, the strategies of applying organoid cultures in translational and clinical studies have not been well developed. In this study, we demonstrate the ability of pancreatic tumor organoids cultures to predict multiple aspects of *in vivo* tumor phenotypes and as a powerful platform for biomarker discovery. In addition to demonstrating that organoids retained genomic alterations of PDX tumors with high accuracy, we report PDO as an *in vivo* relevant model for investigating glycosylation. Lastly, we demonstrate that organoid cultures are an abundant source of tumor enriched extracellular vesicles that can be effectively applied in the clinic as diagnostic biomarkers to identify patients with PDAC from those who have benign diseases such as chronic pancreatitis.

Recent efforts provide strong support for the ability of PDX models to be efficient in predicting drug response in the clinic (1). These results also suggest that PDX models can be an excellent platform for discovering clinically actionable drugs and drug combinations. However, the utility of PDX models in large scale discovery and validation efforts is limited by cost and time. We demonstrate that PDO models are effective in predicting differences in response to therapeutic drugs in PDX models, confirming that PDOs can serve as excellent surrogates for PDX models in drug screening efforts to find new drug combinations that can be subsequently validated in PDX models and translated to the clinic. Our findings highlight an opportunity for exploiting the large collection of patient derived xenograft models available in the scientific community by generating matched sets of PDX and PDO models to accelerate translational research.

Changes in protein N-glycosylation and O-glycosylation can profoundly impact protein maturation, expression, localization, posttranslational modifications and impact its function such as ligand binding and signaling. In addition, aberrant glycosylation could also generate antigens that serve as biomarkers for cancer detection. In fact, the approved pancreatic cancer marker CA19-9 is a glycan antigen. Despite the broad and critical role glycosylation can play, little effort is being placed in understanding glycosylation changes in patient derived models of cancer. We now report that PDO retain glycosylation with high fidelity with those observed in matched PDX models. Increase in abundance of complex type of glycans and fucosylation and sialyation has been associated with cancer (20). Consistently in both PDX and PDO models, complex glycan represents the most frequent type of modification, which is made up of fucosylated and sialyated glycans. Furthermore, we made an unexpected observation that among all the glycans observed, there is a core set of 57 N-glycans that are identified in all patient derived models we analyzed and these 57 glycans collectively represent 50-94% of the relative abundance of N-glycans suggesting that they dominate the glycan landscape in PDAC. Further studies will be needed to confirm this observation in a larger set of samples and to investigate the mechanisms that regulate these glycans in PDAC. Further detailed sequencing of these major glycans requires specialized techniques and are a focus of future studies.

Identification of better blood-based biomarkers for pancreatic cancer diagnosis and disease monitoring is an urgent unmet clinical need. Performing biomarker discovery studies directly from patients’ blood can be limited by high noise effect. We demonstrate a new opportunity to use PDO to discover diagnostic biomarkers by developing a tumor organoid-based pipeline to identify EV proteins from media supernatant with subsequent validation in clinical samples. Previous EV proteomic discovery efforts relied on laborious techniques to isolate EV, typically requiring large amount to supernatant from cancer cell lines (27). We demonstrate that small amounts of media from PDO culture supernatants can be used to enrich for EV allowing for biomarker discovery studies. By combining our discovery efforts with the affinity-based validation platform ExoView, that captures and counts EVs with a target surface protein directly from plasma samples (25), we demonstrate that organoids are a strong biomarker discovery platform with clear clinical utility. In our studies, certain EV markers such as EGFR and HER2 did not clearly distinguish PDAC from chronic pancreatitis patients. While some groups also have not utilized these markers to distinguish PDAC from controls (4, 22), Yang et al demonstrated higher level of these proteins in EVs from PDAC as compared to controls (6). This variance may be due to differences in antibodies used for capturing EVs and distinct detection methods. Yang et al preferentially detected EVs less than 200 nm in diameter while in our study larger EVs were also analyzed. Considering the fact that EVs represent a highly heterogeneous population, different detection methods may capture different subsets of EVs in blood samples. Our ability to identify markers that quantitatively differentiates PDAC and pancreatitis patients highlights the potential of PDOs as a powerful discovery platform for biomarkers and creates new opportunities for discovering context-relevant biomarkers with high fidelity.

Together, our results generate a roadmap for using PDO models as a powerful platform for translational researches aimed at identifying drug sensitivities and new blood-based biomarkers that can be readily applied in the clinic.

## Methods and Materials

### Organoid Culture and Assays

#### Organoid Culture

Organoid cultures were performed as previously described (11). PDX tumors were minced with No.22 blades into 1-2 mm fragments then digested with 1mg/ml collagenase/dispase (Roche) for 30-40 minutes. The digestion was stopped by adding equal volume of 1%BSA in DMEM, then centrifuged at 1500 rpm x 5min. Pellets were further digested with Accutase for 30 minutes then collected by centrifugation at 1500 rpm x 5min. Pellets were then resuspended in organoid growth medium containing Y-27632, 5% matrigel, and growth factors such insulin, FGF2. The suspension was seeded onto 6-well plates pre-coated with matrigel. Culture media was replaced every four days.

#### Drug Treatment Assay

Established organoid cultures were collected and digested as above. For organoids hard to dissociate for single cells, TrypLE was used in place of Accutase. Cells were diluted in organoid growth media at the density of 50,000 cells/ml and 100 ul of the suspension was added into each well of 96 well pre-coated with matrigel. After 4 days of growth, media were replaced with fresh media and drugs were dispensed using a Tecan D300e digital dispenser. Cell death was measured after 4 days using CytoTox Glo (Promega).

#### Morphological and Histological Analysis

Organoids were plated at a density of 25,000 cells/well and images were taken everyday for 12 days. About 200 images were obtained for each line. The images were analyzed for changes using the organoseg software program (13). Briefly, raw images were segmented and analyzed using inbuilt parameters such as area, perimeter and eccentricity. The data were plotted as box plot graph using Prism. To generate organoid tissue sections, they were grown in chamber slides and fixed in 4% PFA for 2 hours followed by incubation with Hematoxylin solution for 10 minutes and washed twice with water. The organoids were scraped and sandwiched between two layers of Histogel (Sigma) using a cryomold and transferred to a tissue cassette followed by fixation in 10% formalin.

### Drug Treatments on PDX Models

#### Establishment of xenografts

Four to six-week-old Foxn1/Nu mice were purchased from Taconic and utilized for these studies. All animal work carried out was approved by the BIDMC Institutional Animal Use and Care Committee (IACUC) and animals were maintained in accordance to guidelines of the American Association of Laboratory Animal Care. To initiate propagation, cryopreserved xenografts were rapidly thawed cut into ~3 × 3 × 3 mm fragments and subsequently implanted subcutaneously in cohorts of 10 mice per PDX line studied, with one small fragments in each mouse. When tumors reached a size of 1500 mm3 they were excised for cohort expansion, cut into ~3 × 3 × 3 mm fragments, and transplanted to the final cohort of mice to be treated with relevant therapeutic agents, with bilateral subcutaneously implantation of fragments in each mouse.

#### Treatment protocol

Xenografts from experimental PDX cohorts were grown to a size of 200-250 mm^3^, at which time mice were randomized and enrolled on study. Dose and schedule of treatments were described in the supplemental materials (**Fig.S3A**). Mice were treated for 28 days and monitored daily for signs of toxicity, with weights and tumor measurements taken three times per week. Tumor length and width was measured using a digital caliper and the tumor volumes estimated using the following formula: tumor volume = [length x width^2^] / 2 (28). Relative tumor growth inhibition (TGI) was calculated by relative tumor growth of treated mice divided by relative tumor growth of control mice (T/C). Experiments were terminated on day 28.

### Glycomic Analysis

#### Preparation of N-glycans from cells or tissue

5×10^6^ cells or 50 mg of tissue sample were used as starting material and were lyophilized. The lyophilized proteins were resuspended in 1 ml of a 2 mg/ml DTT (1,4-Dithiothreitol, Sigma) solution and incubated at 50C for 2h. 1 ml of a 12 mg/ml IAA (Iodoacetamide, Sigma) solution were then added and incubated at RT in the dark for 2h. The DTT and IAA treated proteins were then dialyzed against 50 mM ammonium bicarbonate (Sigma) for 24h at 4C and with the dialysis buffer changed three times. After lyophilization the samples were resuspended in 1 ml of 500 µg/ml TPCK-treated trypsin (Sigma) solution and incubated at 37C overnight. The trypsin reaction was stopped with two drops of 5% acetic acid (Fisherbrand) prior to purification of the digested peptides over C18 200mg Sep-Pak columns (Waters). The Sep-Pak columns were conditioned with 1 column volume (CV) of methanol (Sigma), 1 CV of 5% of acetic acid, 1 CV of 1-propanol (Sigma), and 1 CV of 5% of acetic acid. The trypsin-digested samples were then loaded onto the columns before being washed with 6 ml of 5% acetic acid. Peptides were eluted with 2ml of 20% 1-propanol, then 2 ml of 40% 1-propanol and then 2 ml of 100% 1-propanol. For a given sample, all fractions were pooled and subsequently lyophilized. The lyophilized peptides were resuspended in 200 µl of 50 mM ammonium bicarbonate to which 3 µl of PNGaseF (New England Biolabs) were added for a 4h incubation at 37C. Following this initial incubation another 5 µl of PNGaseF were added for overnight incubation at 37C. The enzymatic reaction was stopped by the addition of two drops of a 5% of acetic acid prior to the purification of the released N-glycans over C18 200 mg Sep-Pak columns conditioned as described above. The PNGaseF-treated samples were loaded onto the columns before being washed with 6 ml of 5% of acetic acid. Flow through and wash fraction containing the released N-gylcans were collected, pooled and lyophilized, and were ready for permethylation. If O-glycans were to be analyzed, the PNGaseF-treated glycopeptides were eluted from the column with 2ml of 20% 1-propanol, then 2 ml of 40% 1-propanol and then 2 ml of 100% 1-propanol. Fractions were pooled and lyophilized, and were ready for O-glycan preparation.

#### Preparation of O-glycans from cells or tissue

400 µl of a sodium borohydride (Sigma-Aldricht) solution ^in 0.1 M NaOH (55mg NaBH^4 ^/ 1 ml 0.1 M NaOH) was added to the lyophilized PGNase F-treated glycopeptides^ and incubated overnight at 45°C. The reactionwasthen stopped by adding drops of pure acetic acid until fizzing stops. The samples were next added to a prepwered Dowex 50W X8 resin (mesh size 200-400, Sigma-Aldricht) column (~3 ml of resin volume) conditioned with 10 ml of 5% acetic acid. The columns were washed with 3 ml of 5% acetic acid. For a given sample, flow through and wash fractions were pooled, collected and lyophilized. The lyophilized samples were next resuspended in 1 ml of an acetic acid:methanol solution (1:9 v/v) and co-evaporated under nitrogen stream. This step was repeated 3 more times. The dried samples were then resuspended in 200 ul of 50% methanol prior to be loaded into C18 200 mg Sep-Pak columns conditioned as above. The columns were washed with 4 ml of 5% acetic acid. Flow through and wash fractions were pooled and lyophilized, and were ready for permethylation.

#### Permethylation of glycans (N- and O-glycans)

Permethylation of N-glycans was carried out to increase sensitivity of MS analysis and performed as follow. Lyophilized N-glycan samples were incubated with 1 ml of a DMSO (Dimethyl Sulfoxide; Sigma)-NaOH (Sigma) slurry solution and 500 µl of methyl iodide (Sigma) for 30 min under vigorous shacking at RT. The reaction was stopped with 1 ml of MiliQ water and 1 ml of Chloroform (Sigma) was added to purify out the permethylated N-glycans. 3 ml of Mili-Q water were then added and the mixture briefly vortexed to wash the chloroform fraction. The water was separated by centrifugation and discarded. This wash step was repeated 3 times and the chloroform fraction was finally dried befor e being redissolved in 200 ml of 50% methanol prior to be loaded into a conditioned (1 CV methanol, 1 CV MiliQ water, 1 CV acetonitrile (Sigma) and 1 CV Mili-Q Water) C18 50 mg Sep-Pak column. The C18 column was washed with 3 ml of 15% acetonitrile and then eluted with 3 ml of 50% acetonitrile. The eluted fraction was lyophilized and then redissolved in 15 µl of 75% methanol from which 1 µl was mixed with 1 µl DHB (2,5-dihydroxybenzoic acid (Sigma)) (5mg/ml in 50% acetonitrile with 0.1% trifluoroacetic acid (Sigma)) and spotted on a MALDI polished steel target plate (Bruker).

#### Data acquisition/analysis

MS data was acquired on a Bruker UltraFlex II MALDI-TOF Mass Spectrometer instrument. Reflective positive mode was used and data recorded between 500 m/z and 6000 m/z for N-glycans and between 0 m/z and 4000 m/z for O-glycans. MS profiles represent the aggregation of at least 20,000 laser shots. Mass peaks were then annotated and assigned to N-/O-glycan composition when a match was found. MS data were further analyzed and processed with mMass (29).

### Analysis of Proteins Markers in Extracellular Vesicles

#### Protein Extraction and Processing

For EV protein extraction, serum-free conditioned media was passed through 0.22 µm filter to remove any floating cells, debris & large extracellular vesicles (> ~250 nm). The media was enriched for secreted vesicles and subject to LC-Ms/MS.

#### LC−MS/MS Analysis & Data analysis

Peptides were analyzed with easy-nLC 1100 (Proxeon, Denmark) coupled to Q-Exactive HF-X. Raw MS files were analyzed by MaxQuant 1.6 with the Andromeda search engine. Tandem mass spectrometry spectra were searched against the “Reference proteome” of human (Taxonomic ID=9606) downloaded from Uniprot. The search included variable modifications of methionine oxidation and N-terminal acetylation, and fixed modification of cysteine carbamidomethylation. Peptides of minimum seven amino acids and maximum of two missed cleavages were allowed for the analysis. False discovery rate of 1% was used for the identification of peptides and proteins. The datasets were then log transformed, quantile normalized and statistically significant changes were determined using empirical Bayes analysis as implemented in the limma package.

#### Human blood collection and processing for extracellular vesicles analysis

Clinical data and blood samples from patients with confirmed histopathologic diagnosis of pancreatic ductal adenocarcinoma, healthy controls and clinical diagnoses of chronic pancreatitis were obtained with IRB approved protocol from May 2018 to September 2018. After obtaining informed consent, whole blood samples were collected in EDTA Vacutainer tubes (BD, Franklin Lakes, NJ). Plasma was obtained after an initial centrifugation of 1,300g x15min. Two additional centrifugations of 2,500g x 15min were performed to remove cellular debris. Remaining plasma samples were stored in aliquots at 80°C.

#### Extracellular vesicle detection by ExoView

Label-free extracellular vesicle (EV) detection was done using a method based on Single Particle Interferometric Reflectance Imaging Sensor (SP-IRIS) which allows multiplexed phenotyping and digital counting of various populations of individual EVs (>50 nm) captured on a microarray-based solid phase chip (25). Capture antibodies against several cancer markers and negative IgG controls were arrayed, in triplicates, on polymer coated sensor chips (30, 31). Organoid culture media containing 0.05% polysorbate 20 were incubated overnight at room temperature on the sensor chips. The unbound particles were then removed by a series of washes: one time in PBS-T (0.05%) and three times in PBS. Each wash was done on a shaker for three minutes. After washing, the chips were incubated at room temperature for 1.5 hours with fluorescently labeled antibodies (CD63 Alexa Fluor 488, CD63 Alexa Fluor 555 and CD9 Alexa Fluor 647) using a solution of 2.5% BSA in PBS-T (0.05%). Following incubation, washes were performed to remove the unbound fluorescently labeled antibody once with PBS-T, three times with PBS and once with Millipore purified water. The chips were then dried. Label-free bound EVs were imaged using the ExoView Reader (NanoView Biosciences, Boston, USA). Analysis was performed using NanoViewer analysis software (NanoView Biosciences, Boston, USA). The effective vesicle binding to the capture antibodies was obtained by subtracting the signals measured after and before sample incubation. Net values from three spot replicates were averaged. Immunostaining of EVs was also detected using the ExoView Reader.

### Data Analysis

#### Whole Exome Sequencing Analysis

Whole exome libraries were prepared using Aglient SureSelect All Exon V5 in accordance with the manufacturer’s protocol. PDXs were sequenced using Illumina HiSeq platform and PDOs were sequenced using Illumina NextSeq platform. The sequencing data were aligned using BWA (32) (v. 0.7.8) to human reference genome, hg19 and variants were identified following the recommended Genome Analysis Tool Kit (GATK) (33) best practice by marking duplicate reads using piccard tools, base recalibration via GATK and variant calling using Unified Genotyper. Since there is no matched normal, the output includes both germline and somatic aberrations. All variants were annotated using Variant Effect Predictor (34)and filtered based on the following criteria: variants with coverage less than 10x and variants with allele frequency > 1% in 1000 Genome project and non-severe variants based on polymorphism, mutation consequences, COSMIC and TCGA mutation frequencies. Concordance of all variants of severe mutation consequences identified in PDX were determine by looking for read evidence for those SNVs in matched PDOs.

#### Drug sensitivity analysis

Organoid cell viability after drug treatment was normalized to mean number of untreated cells. Response to drug concentrations was analyzed using weighted n-parameter logistic regression, ‘nplr’, R-package (35) and the area under the curve (AUC) was estimated using the Simpson’s rule. AUC of each PDO was compared to the TGI of pdx for comparative analysis of drug response.

All <u>*principal component analysis*</u> (PCA) in this article were performed by a singular value decomposition of the normalized and scaled mass-spectrometry data using R-package.

#### Pathway Analysis

Proteins identified in organoid secreted media are analyzed for predictions of protein interactions and their functional associations using STRING database (36) that incorporates known and predicted protein-protein interactions.

## Acknowledgements

We would like to thank members of the Muthuswamy and Hidalgo laboratory for their helpful comments throughout the course of the study. We thank grant or philanthropic support from Judy and Kim Gordon Davies and in part by NIH grant CA224193 to SKM and MH; ERC-2014-ADG-670582 to MH; NIH grant P41GM103694 to RDC; NIH Phase 1 SBIR: 1R43CA213863 to GD and BB; Hirshberg Foundation to LH; Barbara Janson and Arthur Hilsinger Pancreas Fellowship to SDF and Boston University Start up to AE. We would like to thank the patients for their grateful support of the study by providing tissue and blood for research use.

## Author contribution

L.H. conducted experiments, overall experimental design and wrote the manuscript with S.K.M.; B.B prepared the clinical protocol, coordinated obtaining patient samples and reviewed patient information for EV studies; I.P. performed mass spectrometric analysis and data processing for proteomic and phosphoproteomic analysis; D.A. participated in organoid morphology analysis, sample preparation for glycomics, and drug assays; O.G. participated drug assays and coordinated drug treatments on PDX models; A.B. participated in drug assays and sample preparation for proteomic studies; V.S. performed Exoview measurements; E.M. and S.D.L. performed mass spectrometric analysis and data processing for glycomic studies; N.P. and J.G.C. performed drug treatments on PDX mouse models; J.G. prepared clinical protocols for patient samples used in EV studies; R.G. evaluated histopathological features of PDO and PDX; S.P coordinated obtaining clinical information for PDX models; G.D. coordinated experiments on Exoview measurements; M.S.S. and S.D.F helped obtain clinical samples for EV studies; R.D.C. coordinated and contributed experimental design on glycomic study; A.E. coordinated and contributed experimental design on proteomic study; L.B.M performed analysis for omic studies and constructed models for analyzing drug assays; M.H. established PDX models and contributed to experimental design and data analysis; S.K.M. conducted overall experimental design, coordinated collaborations, data analysis and manuscript preparation.

## References

1 Izumchenko E, et al. (2017) Patient-derived xenografts effectively capture responses to oncology therapy in a heterogeneous cohort of patients with solid tumors. Ann Oncol 28(10):2595–2605.

2 Aguirre AJ, et al. (2018) Real-time Genomic Characterization of Advanced Pancreatic Cancer to Enable Precision Medicine. Cancer Discov 8(9):1096–1111.

3 Tiriac H, et al. (2018) Organoid Profiling Identifies Common Responders to Chemotherapy in Pancreatic Cancer. Cancer Discov 8(9):1112–1129.

4 Castillo J, et al. (2018) Surfaceome profiling enables isolation of cancer-specific exosomal cargo in liquid biopsies from pancreatic cancer patients. Ann Oncol 29(1):223–229.

5 Melo SA, et al. (2015) Glypican-1 identifies cancer exosomes and detects early pancreatic cancer. Nature 523(7559):177–182.

6 Yang KS, et al. (2017) Multiparametric plasma EV profiling facilitates diagnosis of pancreatic malignancy. Sci Transl Med 9(391).

7 Anonymous (2015) Essentials of Glycobiology, eds rd, Varki A, Cummings RD, Esko JD, Stanley P, Hart GW, Aebi M, Darvill AG, Kinoshita T, Packer NH, et al.Cold Spring Harbor (NY)).

8 Waddell N, et al. (2015) Whole genomes redefine the mutational landscape of pancreatic cancer. Nature 518(7540):495–501.

9 Cancer Genome Atlas Research Network. Electronic address aadhe & Cancer Genome Atlas Research N (2017) Integrated Genomic Characterization of Pancreatic Ductal Adenocarcinoma. Cancer Cell 32(2):185–203 e113.

10 Oldfield LE, Connor AA, & Gallinger S (2017) Molecular Events in the Natural History of Pancreatic Cancer. Trends Cancer 3(5):336–346.

11 Huang L, et al. (2015) Ductal pancreatic cancer modeling and drug screening using human pluripotent stem cell- and patient-derived tumor organoids. Nat Med 21(11):1364–1371.

12 Kenny PA, et al. (2007) The morphologies of breast cancer cell lines in three-dimensional assays correlate with their profiles of gene expression. Mol Oncol 1(1):84–96.

13 Borten MA, Bajikar SS, Sasaki N, Clevers H, & Janes KA (2018) Automated brightfield morphometry of 3D organoid populations by OrganoSeg. Sci Rep 8(1):5319.

14 Kroep JR, et al. (1999) Gemcitabine and paclitaxel: pharmacokinetic and pharmacodynamic interactions in patients with non-small-cell lung cancer. J Clin Oncol 17(7):2190–2197.

15 Graham MA, et al. (2000) Clinical pharmacokinetics of oxaliplatin: a critical review. Clin Cancer Res 6(4):1205–1218.

16 Yamamoto N, et al. (2012) A Phase I, dose-finding and pharmacokinetic study of olaparib (AZD2281) in Japanese patients with advanced solid tumors. Cancer Sci 103(3):504–509.

17 Flaherty KT, et al. (2012) Phase I, dose-escalation trial of the oral cyclin-dependent kinase 4/6 inhibitor PD 0332991, administered using a 21-day schedule in patients with advanced cancer. Clin Cancer Res 18(2):568–576.

18 Jodrell DI, et al. (2001) 5-fluorouracil steady state pharmacokinetics and outcome in patients receiving protracted venous infusion for advanced colorectal cancer. BrJ Cancer 84(5):600–603.

19 Pinho SS & Reis CA (2015) Glycosylation in cancer: mechanisms and clinical implications. Nat Rev Cancer 15(9):540–555.

20 Holst S, Belo AI, Giovannetti E, van Die I, & Wuhrer M (2017) Profiling of different pancreatic cancer cells used as models for metastatic behaviour shows large variation in their N-glycosylation. Sci Rep 7(1):16623.

21 Almeida A & Kolarich D (2016) The promise of protein glycosylation for personalised medicine. Biochim Biophys Acta 1860(8):1583–1595.

22 Madhavan B, et al. (2015) Combined evaluation of a panel of protein and miRNA serum-exosome biomarkers for pancreatic cancer diagnosis increases sensitivity and specificity. Int J Cancer 136(11):2616–2627.

23 Lewis JM, et al. (2018) Integrated Analysis of Exosomal Protein Biomarkers on Alternating Current Electrokinetic Chips Enables Rapid Detection of Pancreatic Cancer in Patient Blood. ACS Nano 12(4):3311–3320.

24 Osteikoetxea X, et al. (2018) Detection and proteomic characterization of extracellular vesicles in human pancreatic juice. Biochem Biophys Res Commun 499(1):37–43.

25 Daaboul GG, et al. (2016) Digital Detection of Exosomes by Interferometric Imaging. Sci Rep 6:37246.

26 Poruk KE, et al. (2013) The clinical utility of CA 19-9 in pancreatic adenocarcinoma: diagnostic and prognostic updates. Curr Mol Med 13(3):340–351.

27 Brenner AW, Su GH, & Momen-Heravi F (2019) Isolation of Extracellular Vesicles for Cancer Diagnosis and Functional Studies. Methods Mol Biol 1882:229–237.

28 Faustino-Rocha A, et al. (2013) Estimation of rat mammary tumor volume using caliper and ultrasonography measurements. Lab Anim (NY) 42(6):217–224.

29 Strohalm M, Hassman M, Kosata B, & Kodicek M (2008) mMass data miner: an open source alternative for mass spectrometric data analysis. Rapid Commun Mass Spectrom 22(6):905–908.

30 Cretich M, et al. (2011) Silicon biochips for dual label-free and fluorescence detection: Application to protein microarray development. Biosens Bioelectron 26(9):3938–3943.

31 Cretich M, Pirri G, Damin F, Solinas I, & Chiari M (2004) A new polymeric coating for protein microarrays. Anal Biochem 332(1):67–74.

32 Li H & Durbin R (2009) Fast and accurate short read alignment with Burrows-Wheeler transform. Bioinformatics 25(14):1754–1760.

33 McKenna A, et al. (2010) The Genome Analysis Toolkit: a MapReduce framework for analyzing next-generation DNA sequencing data. Genome Res 20(9):1297–1303.

34 McLaren W, et al. (2016) The Ensembl Variant Effect Predictor. Genome Biol 17(1):122.

35 Commo F & Bot BM (2016) nplr: N-Parameter Logistic Regression. https://CRAN.R-project.org/package=nplr.

36 Szklarczyk D, et al. (2017) The STRING database in 2017: quality-controlled protein-protein association networks, made broadly accessible. Nucleic Acids Res 45(D1):D362–D368.

