## supplemental for "Pancreatic tumor organoids for modeling in vivo drug response and discovering clinically-actionable biomarkers"

**A**

| PATIENT | | | | ORIGIN | | | | TREATMENTS | | | |
|---|---|---|---|---|---|---|---|---|---|---|---|
| Xenograft | AGE | GENDER | Ethnic Backgroud | Tumor Type | Tumor AP | Origin | Tumor stage (at tumorgraft collection) | At time of tumorgraft collection | Neoadjuvant | Adjuvant | Mutation Load/Mb |
| Panc-017 | 69 | M | | Pancreas | Adenocarcinoma | Primary Tumor | pT3N0M0 | None | None | None | 12 |
| Panc-030 | 79 | F | Caucasian | Pancreas | Adenocarcinoma | Primary Tumor | Stage IV | None | Xeloda + Radiotherapy | None (exitus 2 months after surgery) | 13 |
| Panc-163 | | | | Pancreas | Ductal Adenocarcinoma | | | | | | 8 |
| Panc-219 | | | | Pancreas | Ductal Adenocarcinoma | | | | | | 16 |
| Panc-265 | | | | Pancreas | Ductal Adenocarcinoma | | | | | | 14 |
| Panc-281 | | | | Pancreas | Ductal Adenocarcinoma | | | | | | 25 |
| Panc-286 | | | | Pancreas | Ductal Adenocarcinoma | | | | | | 39 |
| Panc-014 | 77 | F | Caucasian | Biliar | Extrahepatic biliary adenocarcinoma | Primary Tumor | pT4N1M0 | None | None | None | 10 |

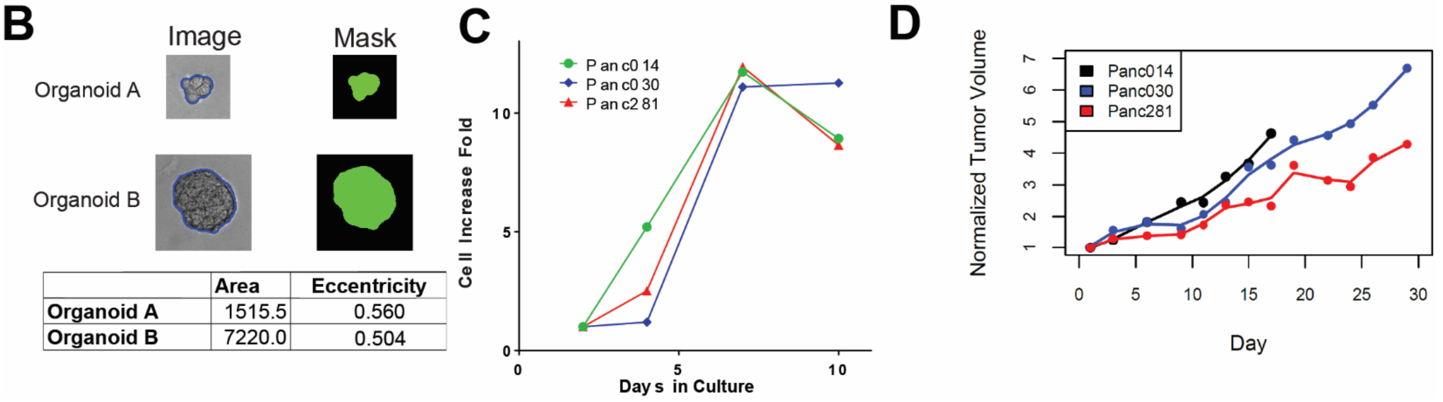


| | Area | Eccentricity |
|---|---|---|
| **Organoid A** | 1515.5 | 0.560 |
| **Organoid B** | 7220.0 | 0.504 |

**Figure S1 Patient derived xenograft models used and properties of the tumor organoid. (A)** Information of patients from whom PDX models were derived. **(B)** Illustration of morphometric analysis on organoids using OrganoSeg. **(C)** Changes of viable cells in organoids cultures from day 2 to day 10. **(E)** Tumor growth in PDX models. Measurements were normalized to initial tumor sizes (200 - 250mm$^3$). mice carrying Panc014 tumors were euthanized early as they reached the humane endpoint as described in IACUC approved animal protocol.

**A**

| Name | Vehicle | Route of Adm | Dose | Total duration of Tx (days) |
|---|---|---|---|---|
| Gemcitabine | 0.9% NaCl | IP injection | 10 mg/kg dose every 4 days | 28 |
| Olaparib | DMSO + PBS | IP injection | 200 mg/kg dose daily | 28 |
| Paclitaxel | Cremophor diluted with 10% Ethanol | IP injection | 30 mg/kg dose every 4 days | 28 |
| 5-Fluouracil | Sterile 5% Dextrose in H2O | IP injection | 100mg/kg dose, once a week | 28 |
| Oxaliplatin | Sterile 5% Dextrose in H2O | IP injection | 10mg/kg dose, once a week | 28 |

**B**

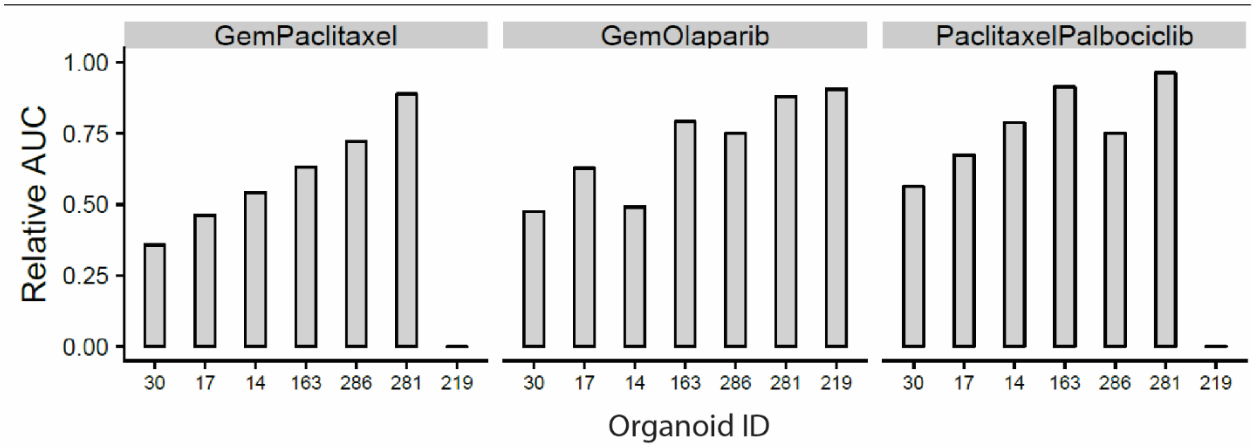


**Figure S2. Drug treatment of patient derived xenograft models (A)** Doses and schedules for treatments on PDX mouse models.  **(B)** Comparison of responses to gemcitabine based treatments and paclitaxel/palbocicilib in organoids.

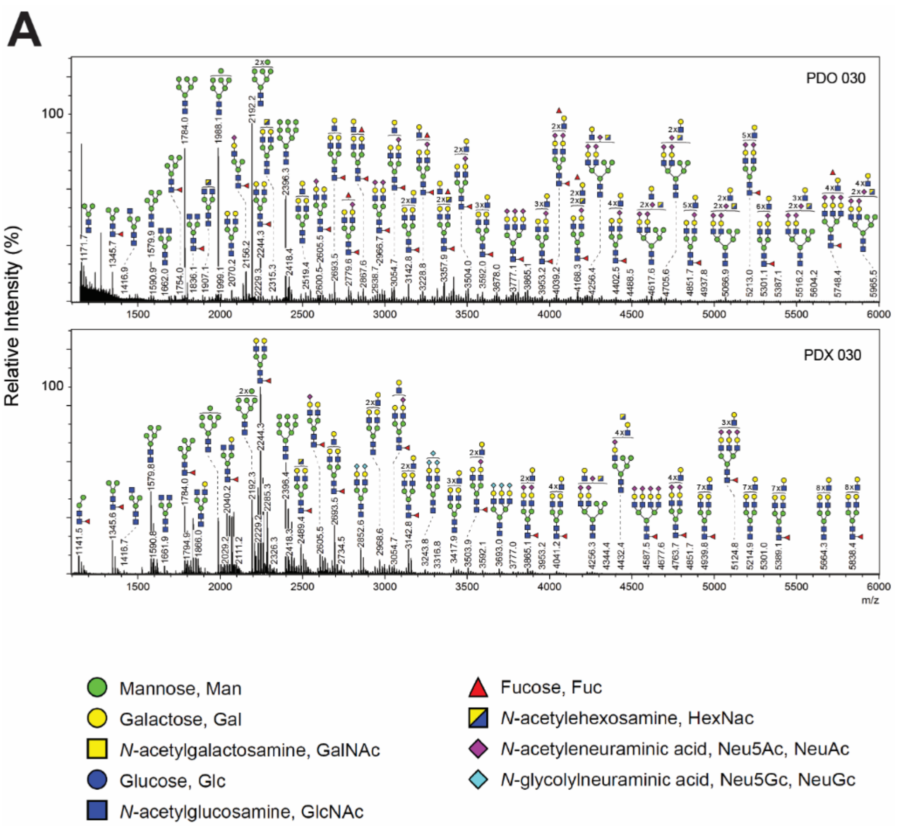


**Figure S3 N-glycan profile of matched PDX and PDO models (A)** Mass spectrometric analysis of N-glycans in PDX and PDO of Panc030. The structures and compositions of glycans were illustrated on top of peaks of corresponding m/z values.

| | Table: Clinical characteristics of subjects enrolled in the extracellular vesicle analysis | | | | | | |
|---|---|---|---|---|---|---|---|
| ID | **Age** | **Gender** | **Disease** | **CA19-9** | **Disease Stage** | **Site of Metastasis** | **Treatment History** |
| PDAC1 | 70 | F | PDAC | 6,707 | IV | Liver | Naïve |
| PDAC2 | 61 | F | PDAC | 175 | IV | Liver | FOLFIRINOX, surgery, radiation, gemcitabine |
| PDAC3 | 83 | M | PDAC | 51 | I | N/A | Naïve |
| PDAC4 | 74 | M | PDAC | 64 | I | N/A | Naïve |
| PDAC5 | 76 | M | PDAC | 264 | II | N/A | Naïve |
| CP1 | 76 | M | Chronic pancreatitis | - | - | - | - |
| CP2 | 36 | F | Chronic pancreatitis | - | - | - | - |
| CP3 | 47 | F | Chronic pancreatitis | - | - | - | - |
| CP4 | 46 | M | Chronic pancreatitis | - | - | - | - |

**Table S1. Information for patients studied in extracellular vesicle measurements**
